# Frailty, polypharmacy, deprescribing and 23-hour activity: insights from a mouse model

**DOI:** 10.64898/2025.12.22.696115

**Authors:** John Mach, Kevin Winardi, Sarah N Hilmer

**Author notes:** **Corresponding author:** John Mach, Laboratory of Ageing and Pharmacology, Kolling Institute, Faculty of Medicine and Health, They University of Sydney and Northern Sydney Local Health District, Sydney, NSW, Australia. **Email:**.

## Abstract

Management of older people with frailty commonly includes polypharmacy and deprescribing. The impacts of frailty on the outcomes of medication use and deprescribing are poorly understood; as are the effects of medication use and deprescribing on frailty and function. Here, using aged C57BL/6J (B6) male mice, we explore the effects of different chronic drug regimens (polypharmacy and monotherapy) and deprescribing on daily activities using an automated behavioral recognition cage, and explore the relationship with frailty and trajectories. At 12 months, male C57BL/6 mice were chronically administered control diet, one of 5 monotherapy diets or one of 3 polypharmacy diets with an increasing Drug Burden Index (measure of total exposure to sedative and anticholinergic medications). At 21 months of age, mice were stratified to continue treatment or to have treatment gradually withdrawn (deprescribed). At 24 months, mice were assessed using the LABORAS automated animal behavioral recognition system for 23 hours. We found that polypharmacy with increasing DBI substantially altered activity and could not be extrapolated from monotherapy response. After deprescribing, while some of the drug effects were reversible, others were irreversible, and we observed some novel changes. Exploring the relationship between frailty and LABORAS behavioral outcomes revealed unique correlations for each intervention group. Four key clusters were identified with different frailty trajectories and attributes, deficits and LABORAS outcomes. This preclinical study demonstrates that medication use and deprescribing can impact activity, frailty and frailty trajectories. The context of polypharmacy and deprescribing are important considerations to enhance translation of pre-clinical studies of frailty.

## Introduction

Older adults internationally experience high prevalences of polypharmacy (taking five or more different medications) and of use of medications with sedative and anticholinergic effects, with greater use in those who are frail (1, 2). Polypharmacy and increasing Drug Burden Index (DBI; a clinical risk assessment tool measuring a person’s total exposure to medications with anticholinergic and sedative effects) (3) are associated with adverse outcomes in older people including functional impairment, falls, frailty, delirium, hospitalisation and mortality (4–6). In addition, medications with anticholinergic effects have been shown to be associated with sleep disturbances and poor sleep quality (7, 8). For this reason, the management of older people with frailty and multimorbidity requires careful assessment of potential benefits and harms of the overall medication burden.

It has been proposed that deprescribing, which is the withdrawal of potentially inappropriate medications supervised by a health care professional with the goal of managing polypharmacy and improving outcomes (9), should remove and may even reverse some of the adverse effects of polypharmacy (10). However, the evidence supporting the benefits of deprescribing on clinical outcomes, including physical and cognitive function, is limited (10).

The lack of high-quality data on the effects of polypharmacy and deprescribing on clinically important outcomes for frail older people is a barrier in developing and implementing optimal strategies to reduce potentially inappropriate polypharmacy (11). Studies on the effects of polypharmacy and increasing DBI on functional outcomes are observational studies, which are likely to be influenced by residual confounding (6, 12). In addition, there are ethical and feasibility challenges in testing the effects of polypharmacy and deprescribing in older frail multimorbid, highly variable adults. To date, some clinical trials of single drugs for a range of conditions have stratified subgroup analyses according to frailty of participants, demonstrating variable effects of frailty on outcomes (13, 14). This has not been applied to testing the effects of frailty on response to polypharmacy. In addition, recent clinical trials of deprescribing have stratified patients according to frailty but have not been powered to detect a prescribing or clinical effect, due to high intra-individual variability and difficulties recruiting frail older people into clinical trials (15). In addition, the dynamic nature of frailty is understudied (16). Frailty trajectories are associated with mortality, cognitive decline, intrinsic capacity and quality of life (16). However, these studies only rarely consider medication impacts (17) on frailty trajectories or transitions.

Advances in preclinical research now enable us to investigate the effect of chronic medication use in monotherapy or polypharmacy, and deprescribing on clinically relevant geriatric outcomes in older frailer individuals (18). In 2021, a preclinical model of chronic polypharmacy and deprescribing was developed, demonstrating that polypharmacy with increasing DBI exacerbated frailty and impaired physical function, which could be reversed with complete deprescribing (19). Automated behaviour recognition systems such as the Laboratory Animal Behaviour Observation Registration and Analysis System (LABORAS) enable collection of activity data over 24-hour light/dark phases. This includes locomotion, immobility, rearing, climbing, grooming, and feeding, which are comparable to measures of physical function and functional independence used in clinical practice, such as activities of daily living. Using this apparatus, age and sex (20), and short-term polypharmacy effects were previously reported (21). There have been no studies investigating the effect of chronic polypharmacy and deprescribing on daily activity over 23 hours and the relationship with frailty. In addition, the dynamics of frailty trajectories are under-investigated in preclinical models.

Here using a clinically relevant aging preclinical model administered chronic polypharmacy and deprescribing we aimed to explore the:

(i) Effects of different chronic drug regimens and deprescribing on behaviour over 23 hours measured using an automated cage (the LABORAS) at age 24 months;
(ii) Association of mouse clinical frailty index with LABORAS outcomes, and
(iii) Trajectories of frailty over time (age 12 – 24 months) to identify different clusters of animals and examine their specific attributes.

## Methods

### Animals

Aging C57BL/6 (B6) male mice, which are commonly used in drug development, preclinical toxicology, and in studies on aging, were the model used for this study. This is a subcohort from the previously described study by Mach et al (19) , using 220 of the healthy animals, sourced and housed at the Kearns Facility, Kolling Institute of Medical Research, Sydney, Australia. A 12-hour light-dark cycle was maintained and animals were housed in boxes of up to five with ad libitum access to food (Rat and Mouse Premium Breeder Diet, Specialty feeds, Western Australia, Australia) and water. This study was approved by the Northern Sydney Local Health District Animal Care Ethics Committee, Sydney, Australia (RESP/15/21).

### Treatment details

At 12 months, as shown in the schematic Figure 1, animals were randomly assigned to treatment groups and were chronically administered control feed (no drugs in food or water), a polypharmacy regimen (five drugs with total zero DBI, low DBI or high DBI in food or water), or a monotherapy feed (simvastatin, metoprolol, oxybutynin or citalopram in food, or oxycodone in water). Medications required to administer the calculated therapeutic dose were mixed with control diet by Specialty Feeds (Western Australia, Australia). Oxycodone (in high DBI polypharmacy regimen and as monotherapy) was administered in the drinking water to enable compliance with regulations for handling and storing of opioids. At 21 months, treatment groups were randomized to continue or withdraw (deprescribe) medication whilst stratifiying on their clinical frailty index score, as previously described (19). At 24 months, animals were tested in the Laboratory Animal Behaviour Observation Registration and Analysis System (LABORAS, Metris, Netherlands).

**Figure 1.**
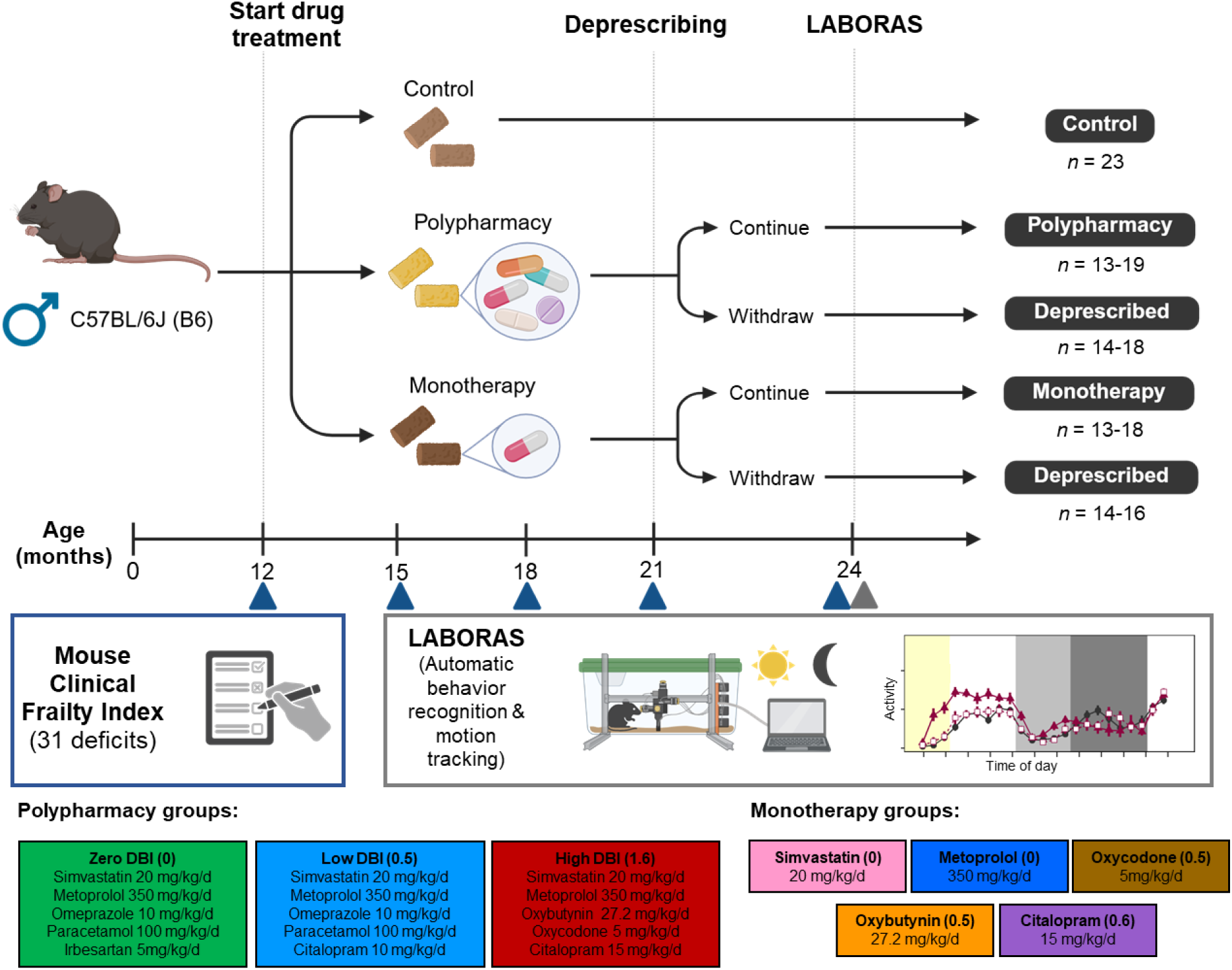
Effect of chronic polypharmacy and monotherapy, and deprescribing on activity measured with the LABORAS: study design. Healthy male C57BL/6J (B6) mice aged 12 months were randomized into 9 treatment groups: control, polypharmacy groups (zero DBI, low DBI and high DBI), and monotherapy groups (oxybutynin, oxycodone, citalopram, simvastatin and metoprolol). Drug Burden Index score shown within brackets beside treatment group. At 21 months of age (after 9 months of treatment), animals on drug treatment were stratified to either continue treatment or to deprescribe, gradually over 6 weeks. At every 3 months from 12 months of age, the mouse clinical frailty index was measured and cross sectionally at 24 months of age, the LABORAS was conducted for 23 hours. Sample sizes are displayed.

The medications for the polypharmacy regimens were selected based on drug classes commonly used by older Australians, which are not likely to be toxic when given alone to a healthy mouse based on previous studies, have similar pharmacokinetics and pharmacodynamics in mice and humans, and are not routinely dose-adjusted in old age.

Therapeutic doses were estimated from previous long-term oral administration of these medicines as monotherapy to mice, based on an observed food intake of 0.11 food/g mouse/ day. The DBI clinical risk assessment tool was used to identify medications to make up the zero, low and high DBI polypharmacy regimens, which have scores of approximately 0, 0.5 (minimum effective dose citalopram) and 1.6 (higher dose citalopram, oxycodone and oxybutynin), respectively. These exposures are consistent with those observed in older adults. For example, amongst community dwelling older Australian men, approximately 75% take medication regimens with zero DBI, 20% low DBI and 5% high DBI (22), while amongst people living in residential aged care facilities approximately 30% take medication regimens with zero DBI, 44% low DBI and 26% high DBI (23).

### Frailty index

As previously described, frailty was measured using the mouse clinical frailty index at 12, 15, 18, 21 and 24 months (19, 24). Briefly, 31 different items relating to multiple body systems (integument, physical/musculoskeletal, vestibulocochlear/auditory, ocular/nasal, digestive/urogenital and respiratory system, and discomfort level and body weight/temperature) were scored 0, 0.5, 1 (except for body weight and temperature) and the overall frailty index score is the sum of all parameters divided by the total number of parameters. The frailty testing was conducted by one experienced observer (J.M.) and such that the tester was blinded to treatment group.

### LABORAS

The LABORAS was used to measure duration spent of 7 different activities (locomotion, climbing, rearing, immobility, eating, drinking and grooming) as well as basic gait parameters (maximum speed, average locomotion speed, and distance travelled), in a 12 hour light/12 hour dark day cycle. In brief, the device was calibrated per manufacturer’s instructions. Food and water were provided *ad libitum*. Animals, food and water were weighed prior to being placed the LABORAS cage, where animals were house individually. As older animals had difficulty reaching the hopper for food, pellets were placed on the cage floor. The LABORAS was set measure from 10:00 am until 09:00 am the next day (23 hours). The 1-hour gap 9:00 - 10:00 was required to clean the cages and organise experiments to run sequentially daily. The LABORAS system detects mechanical signals which was then recorded as hourly duration spent for a given activity. Nonspecific signals were allocated as undefined. The weight of the animals, food, and water were measured before and after, and fecal boli number were counted after each experiment. Up to six animals were tested daily in separate LABORAS cages.

### Data analysis

Data processing, analysis, and visualization was conducted in R/Rstudio (v4.2.0). Data visualization is achieved using ggplot2 (v3.5.2) and ComplexHeatmap (v2.14.0) as appropriate. For LABORAS, mice with complete 23-hour dataset were included in the analysis. Due to technical issues, 18 mice has partial data for the final hour (i.e., total duration <3,600 sec). For those entries, the 23rd hour data was scaled proportionally to represent a full hour. To improve biological reasoning, three mice with complete data were excluded because drinking was not recorded during the 23-hour period. This results in *n* = 220. Exploratory data analysis was conducted using Principal Component Analysis (PCA). Similarly, time was divided into segments based on PCA. For the interest of interpretability in the statistical analysis, the mean time spent for a given activity was calculated within each time segment. Differences between groups were statistically assessed using a two-way mixed ANOVA followed by Tukey *post hoc* test as appropriate whereby the model incorporates both between-(treatment) and within-subject (time segment) design. To assess the association between LABORAS outcomes and frailty index at 24 month, Pearson’s correlation analysis was conducted.

To identify temporal pattern of frailty trajectories across 5 timepoints, clustering analysis was conducted. Optimal number of clusters was determined using NbClust (v3.0.1) package with the following parameters: distance = Euclidean, method = complete, and number of clusters between 3 – 10. Mice with complete frailty dataset across all 5 timepoints were included in the analysis (*n* = 276). Differences in clinical mouse frailty index between the trajectory clusters were assessed with two-way mixed ANOVA followed by tukey *post hoc* test.

Differences in the 31 frailty deficits were compared across trajectory clusters within each month (timepoint) with Fisher’s exact test followed by Bonferroni *p*-value correction. A two-way mixed ANOVA, with treatment included as a covariate, was conducted to assess the association between frailty trajectory clusters and LABORAS outcomes. For all analysis, a *p* < 0.05 was deemed statistical significance.

## Results

### Chronic medications, deprescribing and the LABORAS were well tolerated. Drug exposures altered frailty

We first assessed treatment tolerance and drug intake (Supplementary Figure 1). After 12 months of intervention (12 months chronic treatment or 9 months treatment followed by 3 months deprescribing), compared to control, citalopram significantly increased body weight (Supplementary Figure 1A). Drug intake via food chow, was greater for high DBI polypharmacy treated animals than controls (Supplementary Figure 1C). Compared to control, water intake, body weight and the number of fecal boli were not different for any treatment group (Supplementary Figure 1B, D & E).

### Chronic medication use impacts activities differently, some are reversible by deprescribing

The effect of treatment and deprescribing was examined on LABORAS measured activities (Figure 2 and Supplementary Figure 3-4).

**Figure 2.**
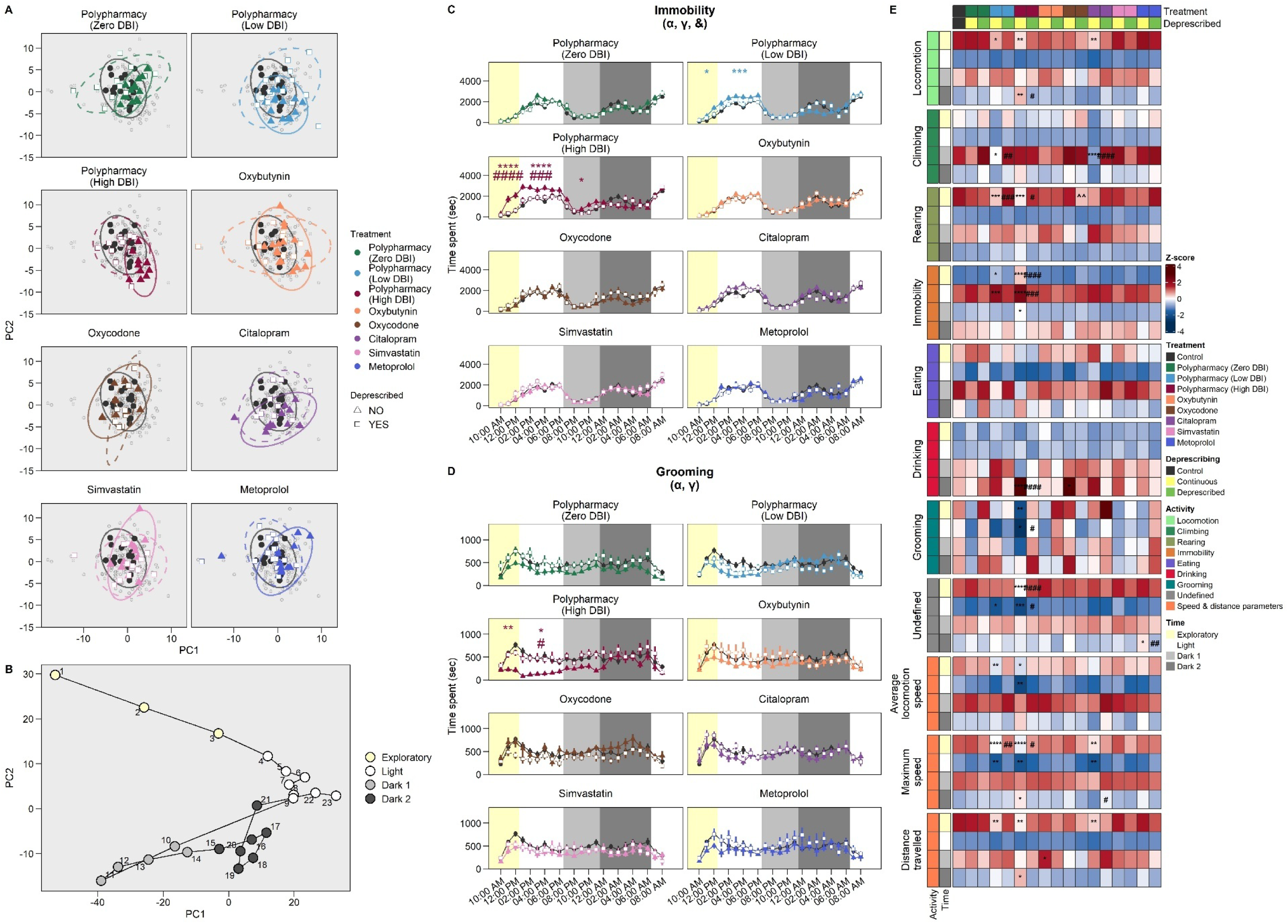
Effect of chronic polypharmacy, monotherapy and deprescribing on LABORAS outcomes. (A) Principal component analysis comparing treatment (filled triangle) and deprescribing (whited square) by applying hourly data for all activities, (B) principal component analysis of temporal shifts in activity patterns following intervention over the course of 23 hours, enabling data-driven time segmentation. Duration spent of (C) immobility and (D) grooming (remaining outcomes shown in Supplementary Figure 3-4). Data displayed as mean ± SEM. Time was segmented into 4 phases. (E) Summary heatmap displaying activities and gait parameters. Cell colors were activity-specific z-scaled, whereby red indicates a longer duration spent whereas blue indicates shorter. Statistical significance was assessed using a two-way mixed ANOVA followed by Tukey *post hoc* test. ANOVA significance is as follow: α: *p* < 0.05 for treatment main effect; γ: time segment main effect; &: treatment × time phase interaction effect. Post hoc results are shown as * is significant comparing continued treatment to control, # is significant comparing continued treatment and the corresponding deprescribing group, and ^ is significant comparing deprescribed group and control. \**p* < 0.05; \*\**p* < 0.01, \*\*\**p* < 0.001; \*\*\*\**p* < 0.0001.

In the PCA plot applying hourly data for all activities (Figure 2A), a tendency toward partial separation was observed between zero, low and high DBI polypharmacy and citalopram from control, which suggest moderate group differences, with some individual samples for each group exhibiting similar characteristics. High DBI polypharmacy, exhibits the most observable partial separation along PC1. Following deprescribing, high DBI polypharmacy deprescribed overlapped with control and is partially separated with continued treatment. All other deprescribed treatment groups overlapped with the control and corresponding treatment group.

To gain deeper insights into temporal shifts in activity patterns following intervention over the course of 23 hours, a PCA plot for time points was constructed (Figure 2B). We identified four distinct time segments that represent the exploratory (first 3 hours; i.e., 10 am – 1 pm), light (1 pm – 7 pm and 7 am – 9 am the following day), dark 1 (7 pm – 12 am) and dark 2 (12 am – 7 am) phases. All activities are unsurprisingly time-dependent irrespective of treatment (Supplementary Figure 2). In the exploratory phase, the mice performed various different exploratory and curious behaviours such as rearing and locomotion. Being nocturnal, they spent majority of the light phase being immobile, indicative of sleeping. Dark phase was divided into 2 parts whereby, in dark 1, mice were generally more active and, in dark 2, they were more immobile. Overall based on definable activities, most time was spent immobile, followed by grooming, locomotion and climbing, with very little time spent rearing, eating and drinking.

Further analysis into each activity demonstrates different chronic medication exposures have differing impacts on activity (Figure 2C-D and Supplementary Figure 3-4). Notably more changes in temporal activity (compared with control) were observed with high DBI polypharmacy, followed by low DBI polypharmacy and citalopram monotherapy. Immobility duration increased in the low and high DBI polypharmacy groups during the first half of the day. Locomotion duration decreased in low DBI, high DBI polypharmacy and citalopram monotherapy groups in the exploratory phase. High DBI polypharmacy was the only treatment to increase locomotion, maximum speed and distance travelled in latter dark phase. For climbing, citalopram and low DBI polypharmacy decreased climbing at the first dark phase. Rearing was reduced by low DBI and high DBI polypharmacy during the exploratory phase. High DBI polypharmacy decreased grooming activity in the exploratory and light phases (Figure 2D). Oxybutynin monotherapy increased distance travelled in the first dark phase. Drinking activity was significantly increased with high DBI polypharmacy and oxycodone monotherapy treatment in dark phase 2. Eating durations did not change with treatment.

Deprescribing reversed some treatment effects (Figure 2C-D and Supplementary Figure 3-4), defined as instances where deprescribing produced changes opposite to those observed when comparing control and treatment group. Notably 50% (9/18) of high DBI polypharmacy temporal activity effects were reversed with deprescribing. This included altered locomotion, immobility, rearing, grooming, drinking, undefined duration and maximum speed. For low DBI polypharmacy, 30% (3/10) of temporal activity effects (decreased climbing, rearing and maximum speed) were reversed with deprescribing. Deprescribing citalopram reversed the complete halt in climbing activity observed with citalopram treatment but did not reverse the decline in locomotion, maximum speed and distance travelled (20%, 1/5). In the exploratory phase, deprescribing oxycodone resulted in a novel (defined as instance where deprescribing produced changes compared to control and no change is observed comparing control and treatment group) decrease in rearing duration. Oxycodone had no treatment effect at this time phase.

### Correlations between LABORAS activities and frailty index varied across treatment groups

We next assessed the association of LABORAS measured outcomes, during different time phases and their total duration over 23 hours, with the mouse clinical frailty index cross-sectionally at age 24 months (19) (Figure 3). Assessing the whole cohort, only weak associations (correlation coefficient < 0.5) with frailty were observed with climbing, maximum speed, average speed during varying time phases.

**Figure 3.**
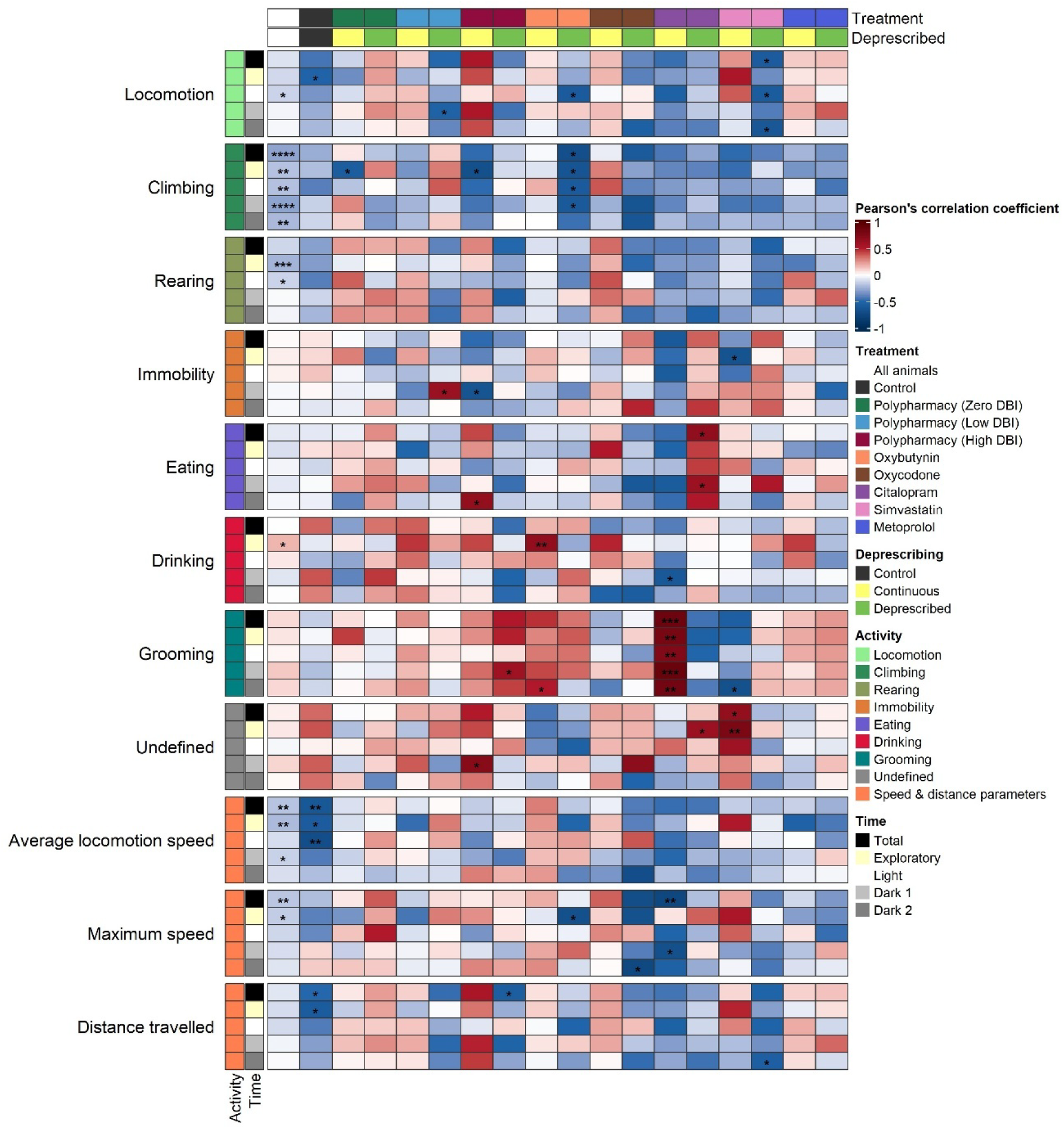
The relationship of LABORAS outcomes over 23 hours with frailty index for treatment and deprescribing groups. Heatmap displaying correlation analysis between LABORAS outcomes and clinical frailty index during the 4 different time segments (row color-coded) or overall (black). Correlation analysis was conducted on each treatment group (column color-coded) or across the whole cohort (i.e., all animals; white). The colour intensity represents the Pearson’s correlation coefficient (r). *p < 0.05; **p < 0.01, ***p < 0.001; ****p < 0.0001.

Further analysis of individual treatments found moderate to strong correlations with frailty, which were very different between treatment groups. For control treated mice only, average speed at the exploratory and light phases and total duration over 23 hours, were moderately correlated with frailty. For High DBI polypharmacy treated mice, climbing (also found for Zero DBI polypharmacy) and immobility were negatively (moderately) correlated with frailty during the exploratory and dark 1 phase, respectively. For citalopram, grooming was strongly positively correlated with frailty at all time phases. No correlation with LABORAS activity was observed for Low DBI polypharmacy, oxycodone and metoprolol monotherapy treated groups.

Similarly, for deprescribed animals, different moderate to high correlations between LABORAS activity and frailty were observed per treatment group. Notably, moderate negative correlations with frailty were found for average speed, max speed, climbing and locomotion for at least one time phase for groups deprescribed from High DBI polypharmacy, oxycodone, oxybutynin and simvastatin, respectively. In the citalopram deprescribed group, frailer mice spent more time eating than those that were less frail, with a moderate positive correlation.

### Frailty trajectories

Using this clinically relevant longitudinal cohort of aged animals that are chronically exposed to medications, we then investigated patterns of frailty trajectories in the whole cohort using the frailty data measured across 5 timepoints (12, 15, 18, 21, and 24 months of age) (Figure 4). Cluster analysis identified 4 key clusters (Figure 4A): (i) progressive increase followed by a decrease between 21 and 24 months (cluster 1), (ii) progressive with a steep slope increase (cluster 2), (iii) progressive with a shallow slope increase (cluster 3), and (iv) a steep increase at middle age and plateau (cluster 4). The trajectory clusters differ in terms of the frailty index at all timepoints, including at 12 months (Figure 4B). The majority of control mice (n = 16/22) are in cluster 3, indicating normal aging trajectory (Figure 4C). Similarly, most of the continuous and deprescribed zero DBI polypharmacy-treated mice (n = 8/15 and n = 10/14, respectively) as well as deprescribed metoprolol monotherapy-treated mice (n = 9/16) are also in this trajectory cluster 3. Cluster 1 is heavily dominated by mice in which treatment was deprescribed, including those deprescribed low DBI polypharmacy (7/15), high DBI polypharmacy (7/13), citalopram (6/11) and simvastatin (7/13). Cluster 2 is not specific to any treatment. Cluster 4, the smallest cluster, is dominated by continuous citalopram monotherapy mice.

**Figure 4.**
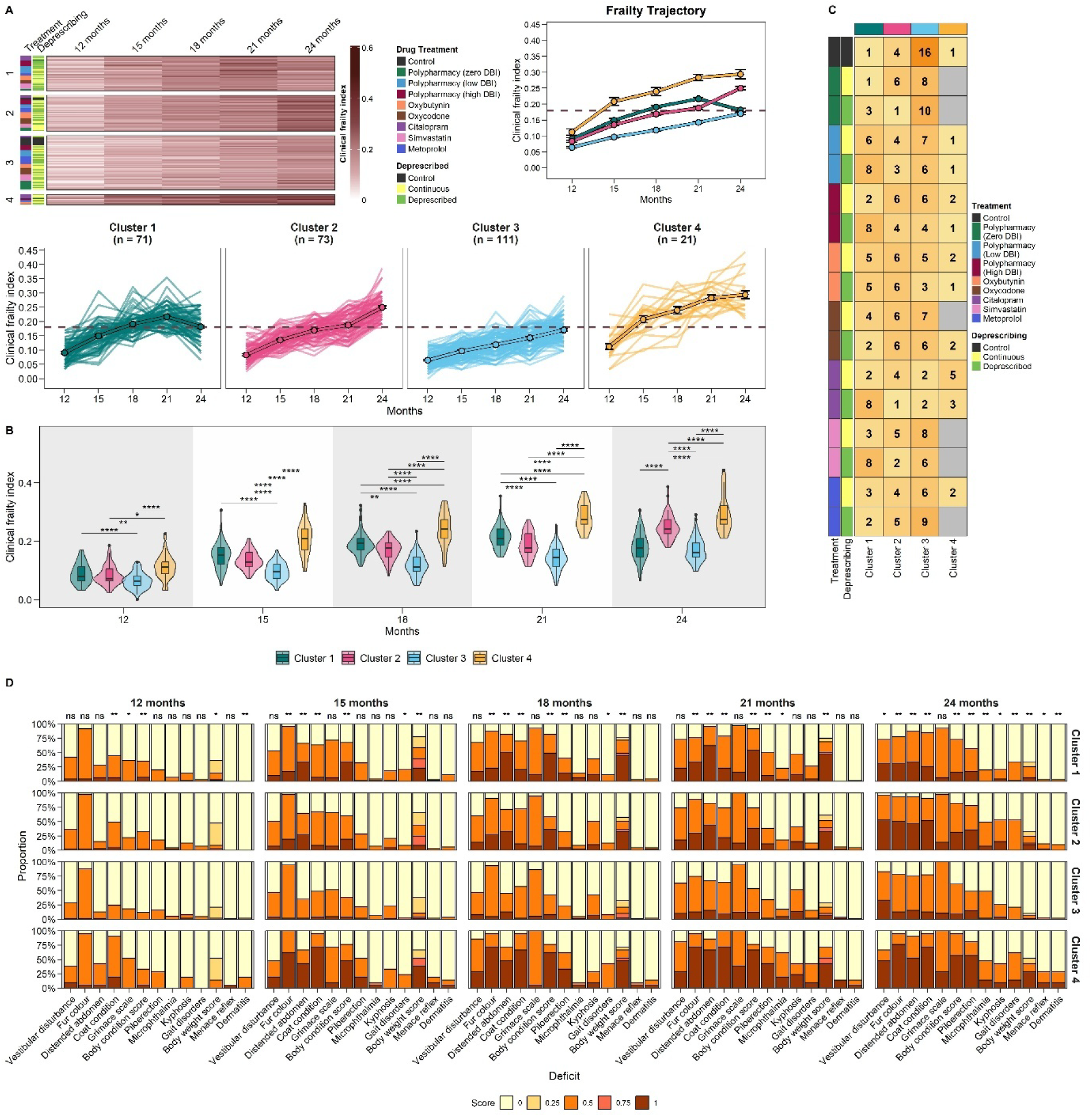
Frailty trajectories and attributes. (A) Cluster analysis showing 4 distinct clusters with different frailty trajectories. Individual-level and population-level (mean ± SEM) frailty scores were depicted at each measured age. (B) Comparing clinical frailty scores between each frailty trajectory cluster at each measured age using a two-way mixed ANOVA followed by Tukey *post hoc* test. Statistical are shown as * between a given cluster pair. (C) Heatmap displaying the sample size of each cluster stratified by drug treatment and deprescribed groups. (D) Stacked bar plots of significantly different frailty deficits between clusters based on Fisher exact test followed by Bonferroni *p*-value adjustment at each time point, where * p<0.05 and ns = non-significant. \**p* < 0.05; \*\**p* < 0.01, \*\*\**p* < 0.001; \*\*\*\**p* < 0.0001.

The individual deficits that make up the clinical frailty index were compared to further explore the differences between frailty trajectory clusters at each measured age (Figure 4D). At 12 months, 5 deficits (grimace scale, body condition, coat condition, body weight and dermatitis) were different between clusters. At 15 months, there were no longer differences in grimace scale and dermatitis across clusters, but body condition, coat condition and body weight remained significantly different, and differences in several other deficits emerged including fur colour, distended abdomen and gait disorders. At 18 months, differences in the same deficits persisted, except for gait disorder, whilst piloerection became different between clusters. At 21 months, the same 6 deficits were different across clusters along with microphthalmia. Finally, at 24 months, 13 deficits differed between clusters: 7 of which were maintained from 21 months (fur colour, distended abdomen, coat condition, body condition, body weight, piloerection and microphthalmia), 2 deficits reemerged from previous timepoints (gait disorders and dermatitis) and differences in 4 new deficits emerged (vestibular disturbance, kyphosis, hearing loss and menance reflex).

Finally, we assessed the differences in LABORAS activity outcomes at age 24 months between frailty trajectory clusters (Figure 5 and Supplementary Figure 6-7). In the exploratory phase, time spent in locomotion and rearing is lower in cluster 2 compared to clusters 1 and 3. Further, we find that in dark phase 1, cluster 1 is more mobile than cluster 3, as measured by duration spent on locomotion, average locomotion speed, and distance travelled. Additionally, average locomotion speed was higher in cluster 1 compared to 3 in dark phase 2. Finally, time spent climbing differed between clusters in dark phase 1: specifically, cluster 3 was longer than clusters 2 and 4, and similarly cluster 1 was longer than cluster 4. All clusters were significantly heavier than cluster 3, however the median body weight for cluster 4 was the greatest.

**Figure 5.**
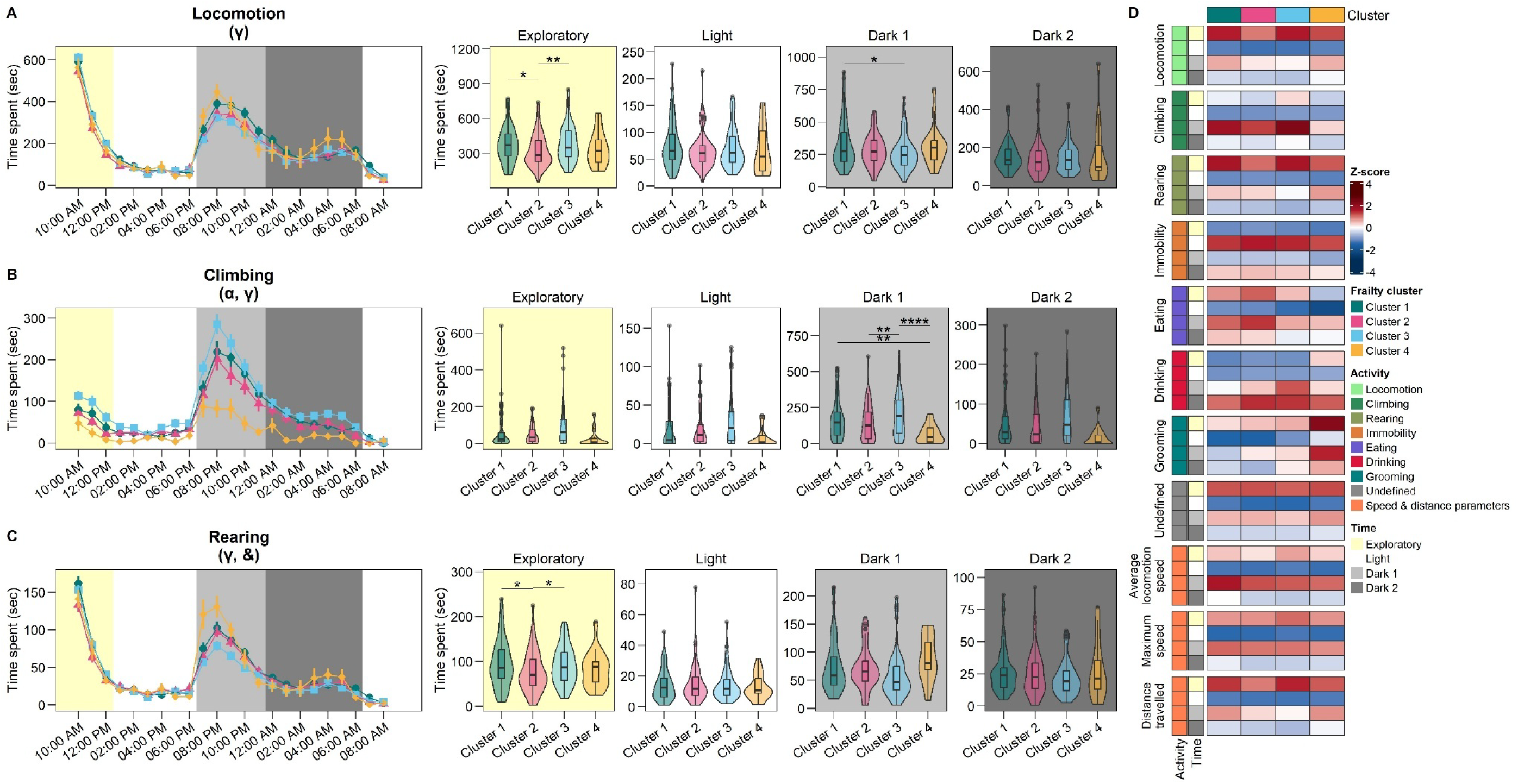
LABORAS outcomes for frailty trajectory clusters. Time spent for (A) locomotion, (B) climbing, and (C) grooming. Left panel depicts hourly data (mean ± SEM) whilst right panel depicts averaged for each time phase (boxplots). Time was segmented into 4 phases. Statistical significance was assessed using a two-way mixed ANOVA followed by Tukey *post hoc* test. ANOVA significance is as follow: α: *p* < 0.05 for cluster main effect; γ: time segment main effect; &: cluster × time phase interaction effect. Post hoc results are shown as * is significant between 2 clusters. \**p* < 0.05; \*\**p* < 0.01, \*\*\**p* < 0.001; \*\*\*\**p* < 0.0001.

## Discussion

This study applied a clinically relevant cohort of aging, polypharmacy and deprescribing, to understand the effects of medication and deprescribing on laboratory mouse activity over 23 hours, using an automated preclinical behavioural recognition system (LABORAS). We explored the relationship between frailty and LABORAS outcomes, which differed across treatment and deprescribing groups. Four clusters of mice were identified with different frailty trajectories from middle to old age, which differed in medication exposures, frailty deficits and LABORAS activities. Polypharmacy is common in old age, and interventions on frailty should consider testing in the context of polypharmacy to enhance translation.

Consistent with clinical observational studies, we found that polypharmacy with increasing DBI impairs physical activity (locomotion, immobility, climbing, rearing) and activities of daily living (grooming). Interestingly, medication regimens; particularly low, high DBI polypharmacy and citalopram monotherapy, with greater frailty had consistently greater number of altered activities compared to control. The effect of high DBI polypharmacy treatment could not be extrapolated by monotherapies as previously observed with open field based gait analysis (25) and hepatic proteome changes (26). For monotherapy, only citalopram, oxybutynin and metoprolol treatment altered definable activity. In addition, changes in activity were not observed for all polypharmacy groups, however a step-wise increase was observed with polypharmacy with increasing DBI score, consistent with polypharmacy’s effect on gait (25). For high DBI polypharmacy treated animals only, immobility was observed in the exploratory and light phase, which could be due to the anticholinergic and sedative medications in the regimen. However increased activity in the latter dark phase, which could be reversed by deprescribing. The latter dark phase represents a time when rodents is generally when go back to sleep and the hyperactivity shows close resemblance to sundowning syndrome or potentially disturbed sleep found in patients that can be triggered by certain medication regimens. In addition, in humans increased anticholinergic burden have been shown to be associated with sleep disturbances and poor sleep quality (7, 8). Together the results suggest, medications particularly in polypharmacy have complex effects. This research supports the need for present and novel therapeutics to be tested in preclinical models in the context of medications they generally are used in, by older frail people.

Deprescribing was observed to reverse some of the medication effects in the groups most affected; namely high DBI, followed by low DBI polypharmacy. This is consistent with our previous open field gait analysis study (25). This included reversing drug effects on mobility (locomotion, immobility, maximum gait speed, rearing, climbing), and personal hygiene and activities of daily living (grooming). Interestingly, in this study we find low DBI polypharmacy had a lower proportion of reversible deprescribing effects than high DBI polypharmacy. Contrary to the previously reported with computational open field gait analysis findings (25), where increasing DBI polypharmacy regimens had less deprescribing reversibility. For monotherapies, deprescribing citalopram restored the climbing activity in the first dark phase. We also observed a decrease in rearing in oxycodone deprescribed animals, which was unique to deprescribing and not an oxycodone treatment effect. These results together, demonstrate that deprescribing effects are complex; can be reversible, irreversible or cause novel changes. Deprescribing inappropriate prescribed medication can reverse some of the medication-related activities, consistent with our gait analysis (25).

However, reversibility may not be proportional to the number of activity changes or DBI score. In addition, different measurements and sensitivity of test can pick up different number of deficits and reversibility can vary. Notably, as observed in clinical studies, deprescribing did not consistently yield benefits, but no further harm was detected (27). Together our results suggest deprescribing does not cause harm and may be beneficial.

The effect of medications on frailty is poorly understood due to limited studies and complex due to the wide variety of drugs, combinations and indications. Preclinical studies investigating frailty commonly look at healthy ageing rodents and do not generally consider medication use which is common with older people to manage multimorbidity. Several studies investigating geroprotective agents have reported reversals in frailty (28–31), however their benefit in the context in older individuals taking polypharmacy is yet to be evaluated.

Notably, the only preclinical study investigating whether chronic polypharmacy causes frailty demonstrated that chronic polypharmacy increased frailty, which as attenuated or reversed with deprescribing (19). Exploring the relationship between frailty and LABORAS outcomes, we found unique correlations for each treatment and deprescribing group. Importantly for the whole cohort and healthy ageing controls, increasing frailty is associated with decreased average gait speed, consistent with clinical studies (32–34). However, when we explored interventions; polypharmacy, monotherapy and deprescribing, other distinct frailty correlations were found suggesting that the relationship between frailty score and activity outcomes is complex and can be impacted by medication use and deprescribing.

A growing number of clinical studies have shown that individuals can have different frailty trajectories and increasing frailty trajectories are associated with higher risk of adverse outcomes such as emergency department visits, hospitalization, and all-cause mortality (35–39). However, these studies rarely consider medication impacts. Using this clinically relevant preclinical cohort, which applied common medication exposures in old age, we explored the different frailty trajectories to further understand the dynamic nature of frailty. We identify 4 key clusters with different frailty trajectories and characteristics similar to clinical studies.

Cluster 3 the ‘healthiest aging’ cluster, had the lowest baseline frailty score, shallowest slope increase and was characterized with lowest body weight, climbed more than cluster 2 and 4 in Dark 1 period and was predominantly made up of control animals (72%). Cluster 4 could represent a ‘medication-related obesity’ group, it had the steepest increase followed by a plateau, had the greatest number of deficits, mice were generally heavier, climbed the least in Dark 1 period, and the group was dominated by continuous citalopram monotherapy animals (24%). Interestingly consistent with this study, a clinical prospective cohort study found that worse trajectories were associated with obesity (38). Finally, Clusters 1 and 2 were similar.

Cluster 2 can be described as ‘general frailty with medications’ with no specificity to any treatment. Cluster 1 is heavily dominated by mice that underwent deprescribing, respresenting cluster whereby ‘frailty improves following deprescribing’. Cluster 1 is characterised by more locomotion and rearing in the exploratory phase than cluster 2, demonstrating deprescribing can have beneficial effects on activity and frailty for some individuals. Collectively, we found frailty clusters showing similiar trajectories and attributes to clinical studies. This preclinical model demonstrated the dynamic nature of frailty showing how individual variations, medication and deprescribing can alter trajectories. Future research is required to examine the mechanisms behind the different clusters, and confirm the impact of medications and deprescribing on frailty trajectories and activity in humans

The strength of this study is that the outcomes were measured using an internationally validated method that automatically, unbiased and objectively measures activities of lab murine over light/ dark phases, particularly capturing the nocturnal behaviour of rodents in the dark phase. The measured activities are comparable to automatically detecting activities daily living from wearable gadgets (40) and in-home sensors in humans (41), such as transferring/ mobility (locomotion, immobility, rearing, climbing), personal hygiene and dressing (grooming), and feeding (eating and drinking). The medication exposures investigated are highly clinically relevant, as they mimic chronic use of medications, as well as the context that they would be used, polypharmacy, in older people. In addition, as observed in clinical observational studies, the polypharmacy mouse model consistently demonstrates that polypharmacy with increasing DBI is associated with functional decline and impairment in activities of daily living (6).

There are some limitations in this study that guide interpretation and could be addressed with future research. The mice in this study had no pre-existing morbidities for which the medication was prescribed, unlike medication use in humans, limiting the model unable to explore medication-disease interactions. It is possible that medication-disease interactions could impact the trajectory of frailty differently if the drugs effectively managed disease.

Future studies are required to confirm these findings in patients with pre-existing illnesses. In addition, examining deprescribing individual medications in the context of polypharmacy was not examined. This study focused on male mice and future studies should confirm these findings in females, especially since older women have a higher prevalence of polypharmacy (42, 43) and there are known sex differences in response to polypharmacy in pre-clinical (44–46) and clinical studies(47). In addition, the results reported here differed between medication groups and may be specific to the medicines and doses selected. Future studies are required to elucidate the effects of polypharmacy regimens containing varying classes, combinations and doses of medicines. We could not control the medication intake in this study, as the feeding was *ad libitum*. In addition, low DBI and high DBI polypharmacy food tended to easily crumble reducing the accuracy of weighing the food to calculate the food and drug intake.

The exploration phase affects the regular daily activities of the mice during the light phase. In the future, undisturbed recording over 2 days can attain results after acclimatization over 24 hours.

## Conclusion

In conclusion, for the first time using an automated animal behaviour recognition device, we demonstrate that chronic polypharmacy with increasing DBI impairs mice daily activities.

Our results also show that deprescribing is not harmful and can partially reverse treatment effects on old aged C57BL/6 male mice. We demonstrated that treatment and deprescribing have distinct associations with frailty that are different to non-treated animals. We further identify clusters that display different frailty trajectories and attributes, consistent with some clinical studies. Collectively, our results support the inclusion of polypharmacy regimens in studies of frailty and activity to enhance translational value of the research, as they are key elements of the real-world clinical setting of older people with multimorbidity.

## Acknowledgments

We thank Dr Chris Little for his advice on the use of the LABORAS. The authors acknowledge laboratory assistance from Gizem Gemikonakli, Caitlin Logan, Swathi Ekambareshwar, Doug Drak and Jennifer Debenham. The authors also acknowledge the support of the Kearns facility staff, Kolling Institute. The graphical abstract and experimental design image (Figure 1) was created with https://www.BioRender.com/. Open access publishing facilitated by The University of Sydney, as part of the Wiley - The University of Sydney agreement via the Council of Australian University Librarians.

## Author contribution

S.N.H., J.M., and K.W. conceptualized and conceived the project. J.M. conducted and coordinated acknowledge research staff to conduct the LABORAS experiments and animal wellfare checks. J.M. processed the LABORAS data. J.M. conducted all clinical frailty index experiments. K.W. collated all datasets, designed the analytical framework, and conducted all associated statistical analysis and data visualization. J.M. and K.W. wrote the original draft of the paper with oversight and direction from S.N.H. All authors provided the critical feedback and/or edited the mansucript and approved the final version for submission.

---

**Supplementary Figure 1.**
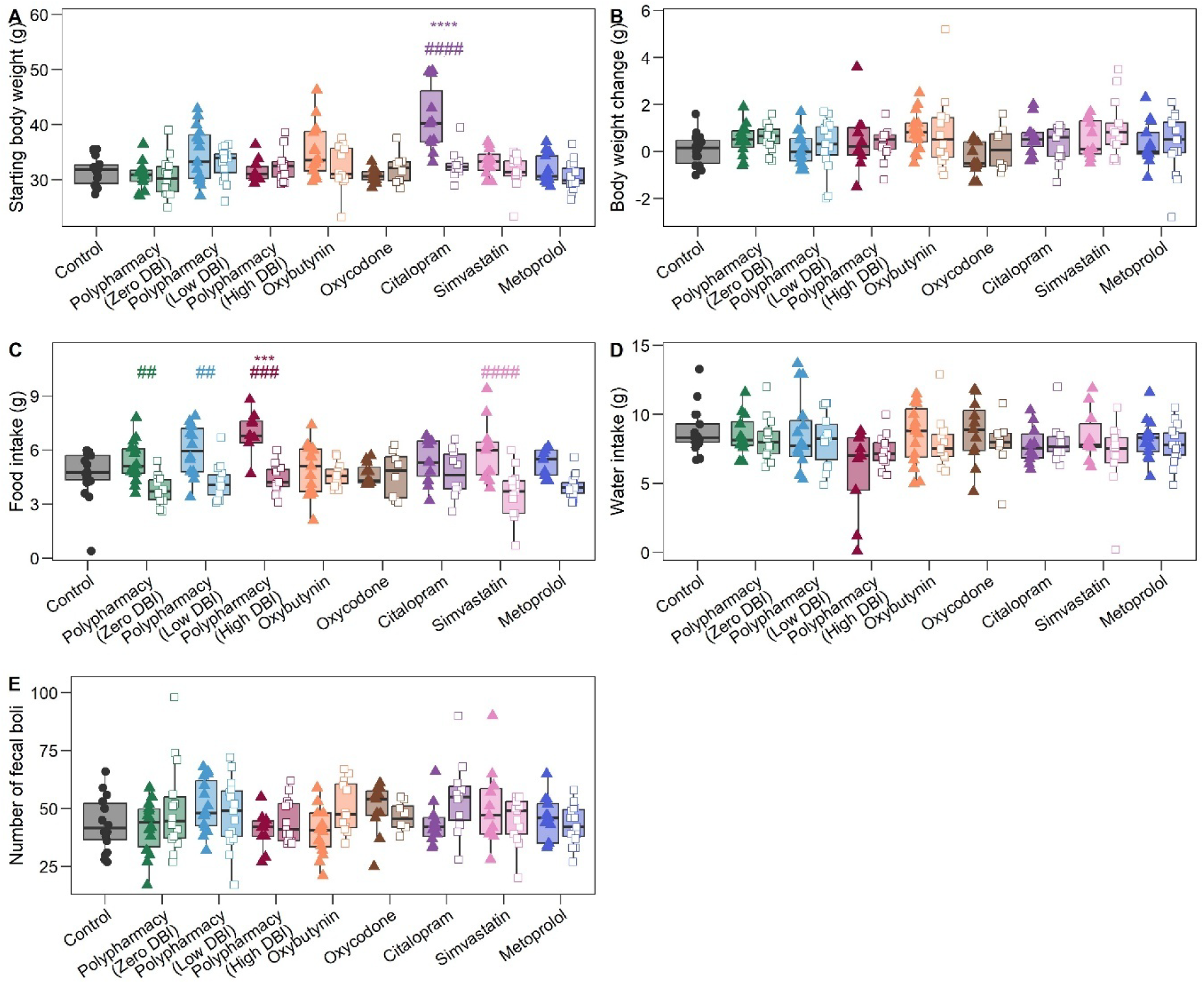
Effect of chronic polypharmacy and monotherapy, and deprescribing on wellbeing before and after LABORAS. (A) Starting body weight before LABORAS, and (B) body weight change, (C) food intake, (D) water intake, and (E) number of fecal boli after LABORAS. Statistically significant was determined with one-way ANOVA followed by a Tukey *post hoc* test. Difference shown as * is significant compared to control, # is significant comparing continued treatment and the corresponding deprescribing group. \**p* < 0.05; \*\**p* < 0.01, \*\*\**p* < 0.001; \*\*\*\**p* < 0.0001

**Supplementary Figure 2.**
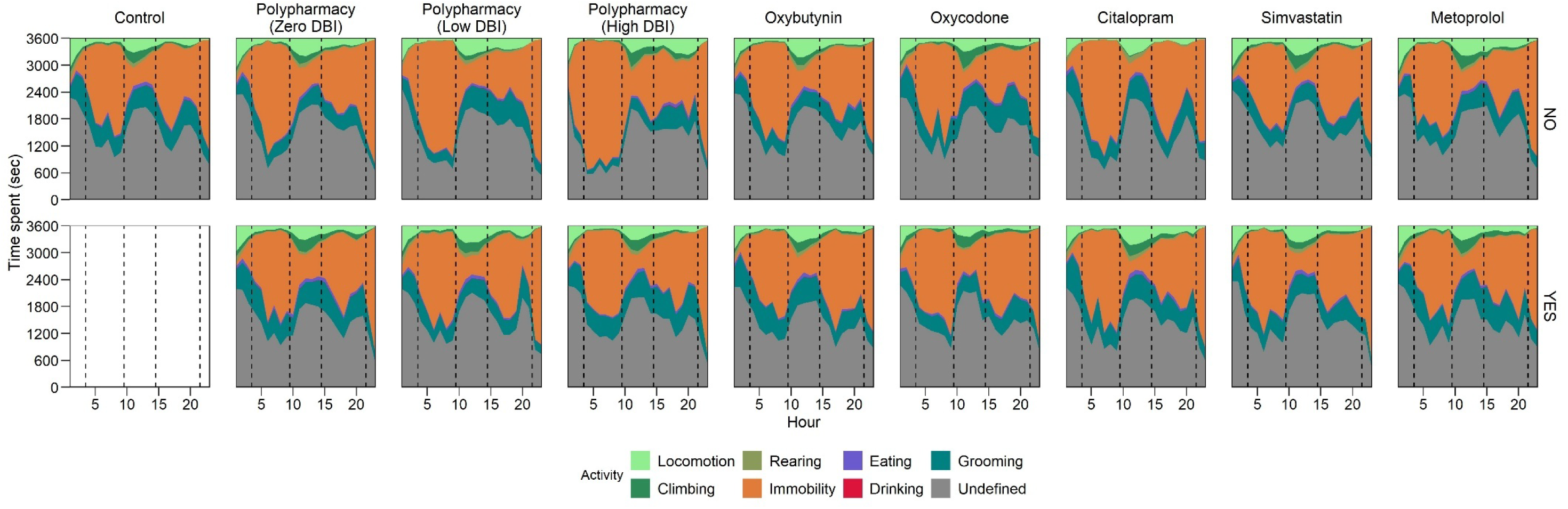
Area chart showing activity duration patterns over time with continued treatment (NO) on the upper panel and the corresponding deprescribed (YES) on the lower panel. Dotted lines separate the 23 hours into the time phases.

**Supplementary Figure 3.**
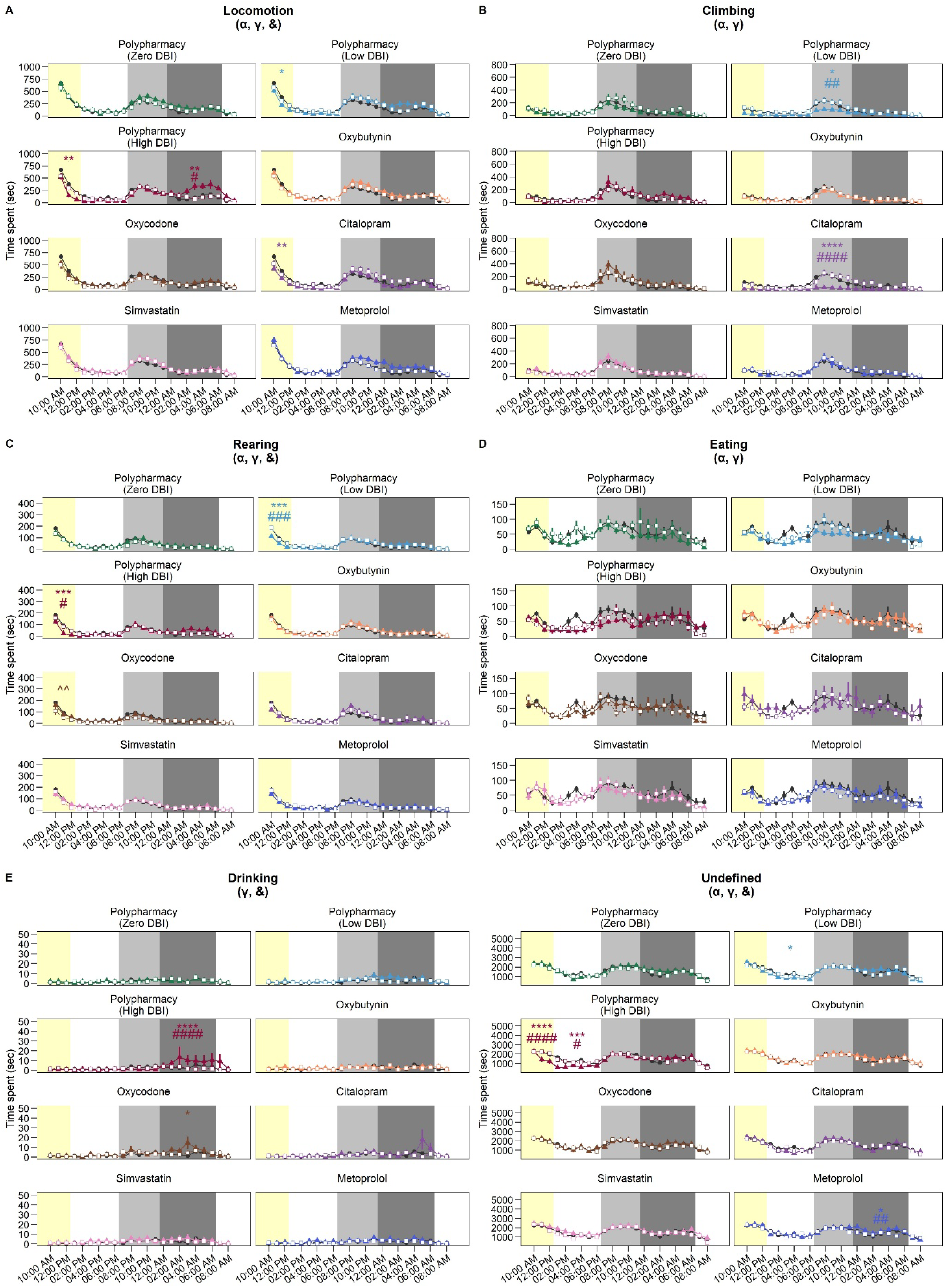
Effect of chronic polypharmacy and monotherapy, and deprescribing on LABORAS outcomes. Duration of the LABORAS-measured activities incuding (A) locomotion, (B) climbing, (C) rearing, (D) eating, (E) drinking and (F) undefined. Data displayed as mean ± SEM. Time was segmented into 4 phases. Statistical significance was assessed using a two-way mixed ANOVA followed by Tukey *post hoc* test. ANOVA significance is as follow: α: *p* < 0.05 for treatment main effect; γ: time segment main effect; &: treatment × time phase interaction effect. Post hoc results are shown as * is significant compared to control, # is significant comparing continued treatment and deprescribing group, and ^ is significant comparing deprescribed group and control. \**p* < 0.05; \*\**p* < 0.01, \*\*\**p* < 0.001; \*\*\*\**p* < 0.0001.

**Supplementary Figure 4.**
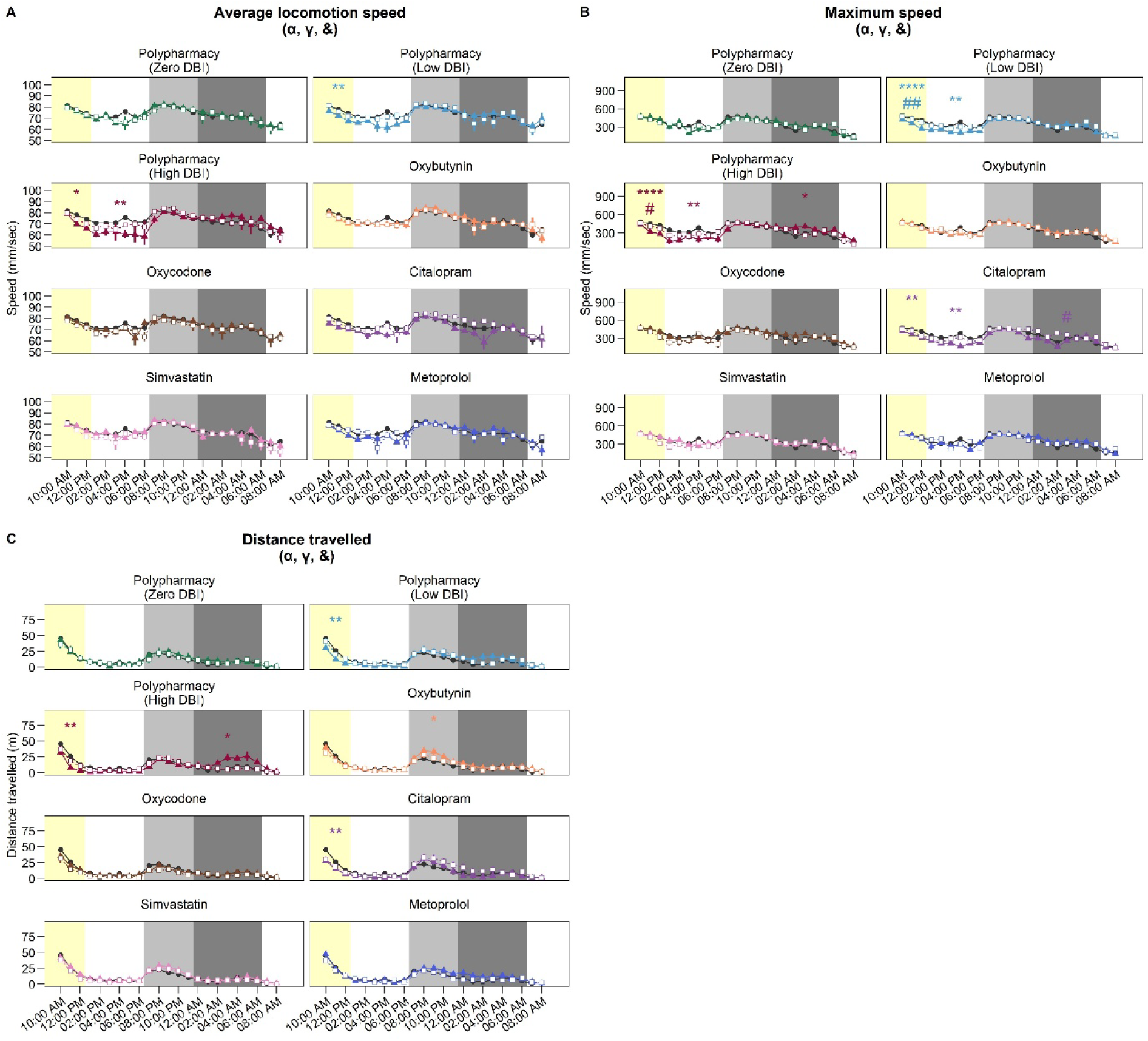
Effect of chronic polypharmacy and monotherapy, and deprescribing on LABORAS gait outcomes. LABORAS-measured gait parameters incuding (A) average locomotion speed, (B) maximum speed, and (C) distance travelled. Data displayed as mean ± SEM. Time was segmented into 4 phases. Statistical significance was assessed using a two-way mixed ANOVA followed by Tukey *post hoc* test. ANOVA significance is as follow: α: *p* < 0.05 for treatment main effect; γ: time segment main effect; &: treatment × time phase interaction effect. Post hoc results are shown as * is significant compared to control, # is significant comparing continued treatment and deprescribing group, and ^ is significant comparing deprescribed group and control. \**p* < 0.05; \*\**p* < 0.01, \*\*\**p* < 0.001; \*\*\*\**p* < 0.0001.

**Supplementary Figure 5.**
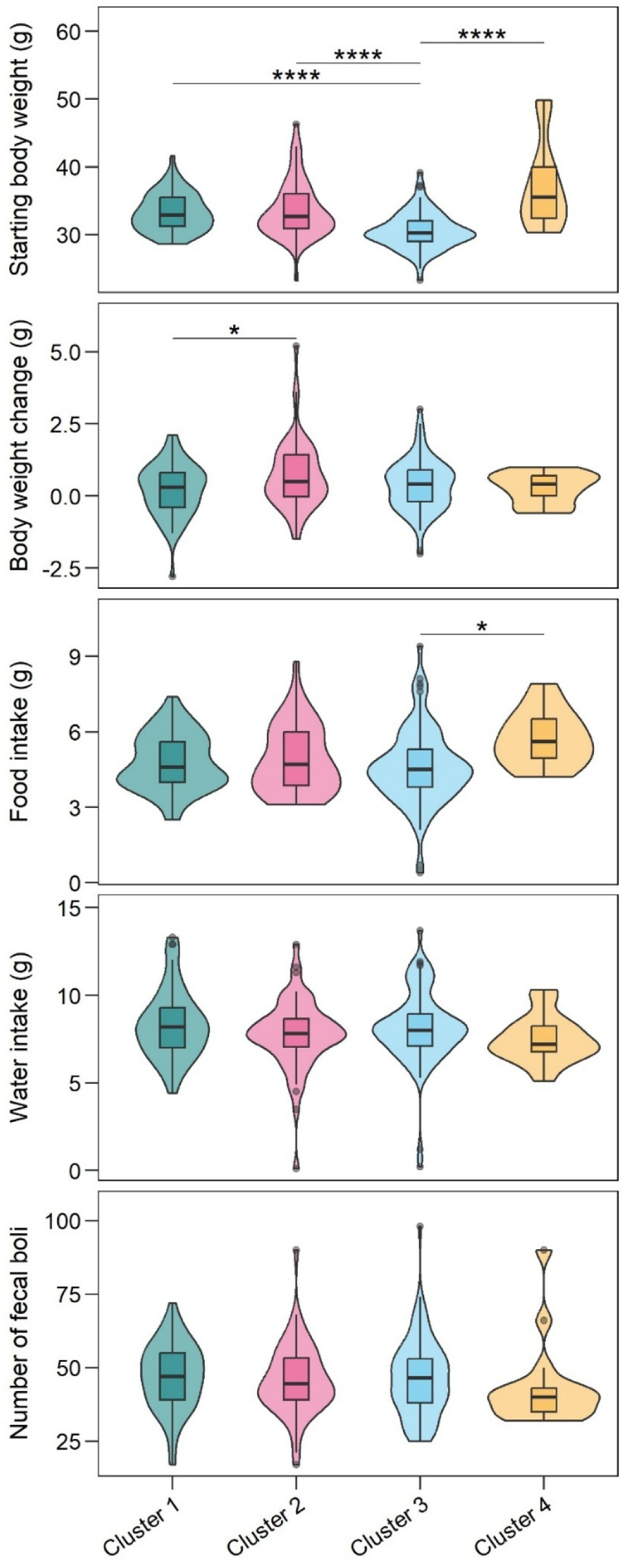
Effect of frailty trajectory on wellbeing before and after LABORAS. (A) Starting body weight before LABORAS, and (B) body weight change, (C) food intake, (D) water intake, and (E) number of fecal boli after LABORAS. Statistically significant was determined with one-way ANOVA followed by a Tukey *post hoc* test. Difference shown as * is significant between frailty trajectory clusters. \**p* < 0.05; \*\**p* < 0.01, \*\*\**p* < 0.001; \*\*\*\**p* < 0.0001.

**Supplementary Figure 6.**
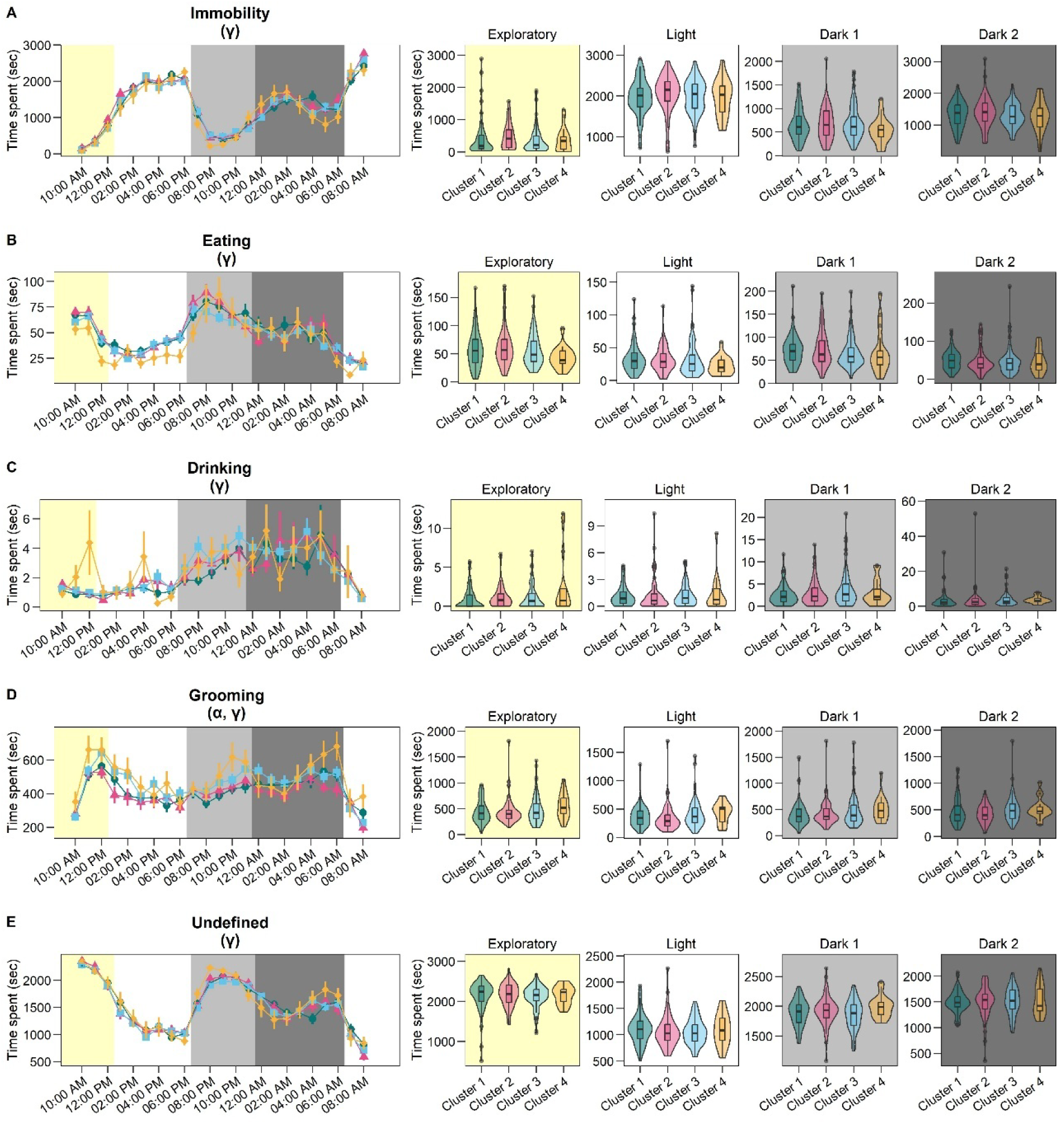
Effect of frailty trajectory cluster on LABORAS outcomes. Duration spent for (A) immobility, (B) eating, (C) drinking, (D) grooming, and (E) undefined. Left panel depicts hourly data (mean ± SEM) whilst right panel depicts averaged for each time phase (boxplots). Time was segmented into 4 phases. Statistical significance was assessed using a two-way mixed ANOVA followed by Tukey *post hoc* test. ANOVA significance is as follow: α: *p* < 0.05 for cluster main effect; γ: time segment main effect; &: cluster × time phase interaction effect. Post hoc results are shown as * is significant between 2 clusters. \**p* < 0.05; \*\**p* < 0.01, \*\*\**p* < 0.001; \*\*\*\**p* < 0.0001.

**Supplementary Figure 7.**
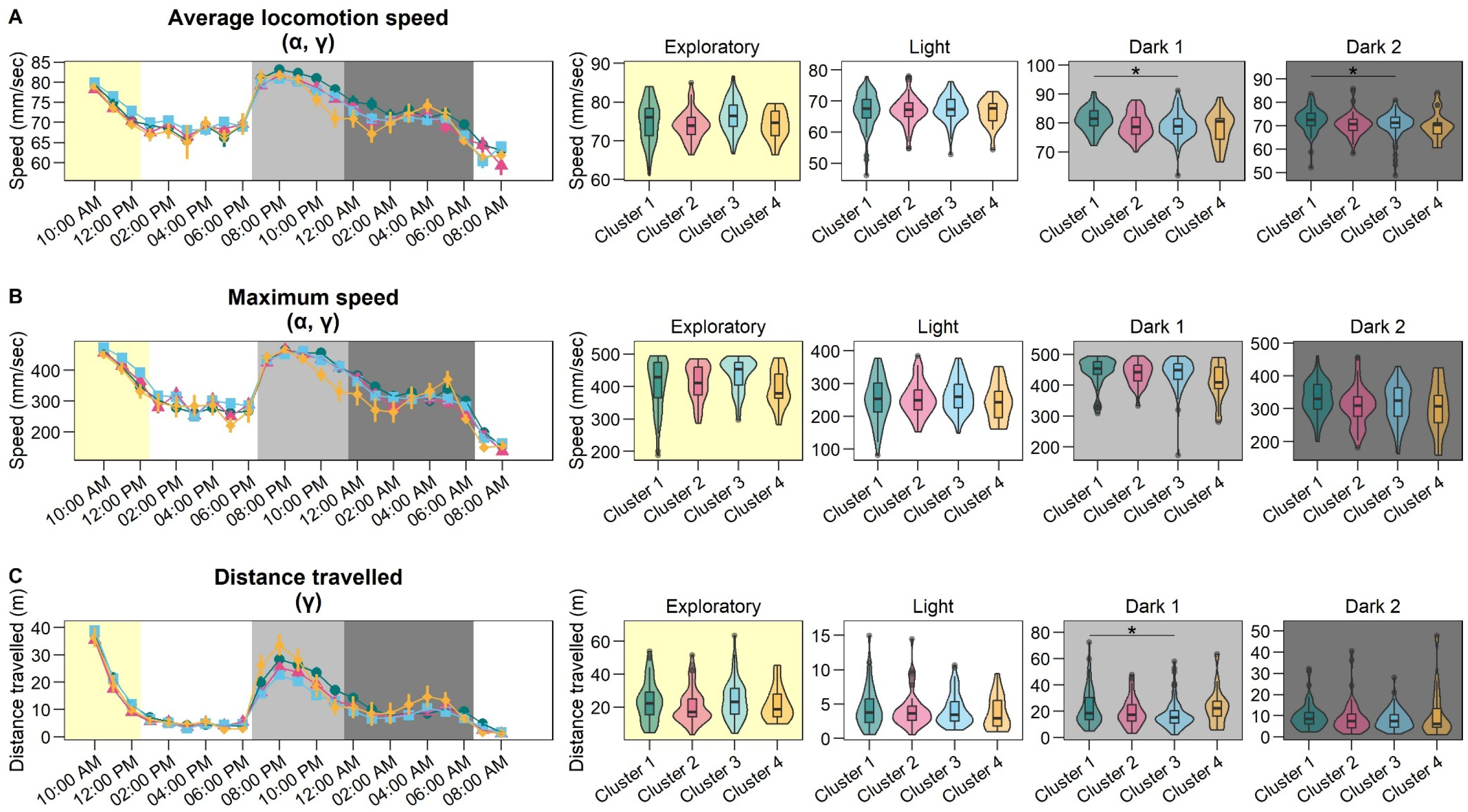
Effect of frailty trajectory cluster on LABORAS gait outcomes. LABORAS-measured gait parameters incuding (A) average locomotion speed, (B) maximum speed, and (C) distance travelled. Left panel depicts hourly data (mean ± SEM) whilst right panel depicts averaged for each time phase (boxplots). Time was segmented into 4 phases. Statistical significance was assessed using a two-way mixed ANOVA followed by Tukey *post hoc* test. ANOVA significance is as follow: α: *p* < 0.05 for cluster main effect; γ: time segment main effect; &: cluster × time phase interaction effect. Post hoc results are shown as * is significant between 2 clusters. \**p* < 0.05; \*\**p* < 0.01, \*\*\**p* < 0.001; \*\*\*\**p* < 0.0001.

